# New relationships are not necessarily more variable: female degus are more consistent with strangers, male degus are less

**DOI:** 10.1101/2021.08.19.456980

**Authors:** Amber Thatcher, Nathan Insel

**Affiliations:** Department of Psychology, University of Montana, Missoula, Montana, USA

**Keywords:** degu, exploration, learning, peer, relationship, social

## Abstract

When an organism explores a new environment or stimulus it varies its behavior to ensure proper sampling. As contingencies are learned, variance can give-way to routines and stereotypies. This phenomenon is common across species but has not been well studied in the social domain, in which the stimulus an animal investigates, another individual, may react negatively to unexpected behaviors. Here we investigate the effects of social familiarity on interaction variability in degus, female members of which are predisposed to form relationships with new, same-sex individuals. Degus were presented with a series of 20 minute, dyadic “reunion” sessions across days, interleaving exposures to familiar and unfamiliar same-sex conspecifics. We found that session-to-session variability in males was higher between strangers compared with cagemates, suggesting males may establish relationships by testing different social roles. In contrast, following an initial exposure, female strangers showed lower session-to-session change compared with cagemates, potentially establishing new relationships by maintaining behavioral norms. Social novelty did not appear to affect variability of interaction timing within a session. Given ecological pressures on female degus to form large, stable social networks, the data are consistent with the notion that higher behavioral variability across encounters is maladaptive for establishing cooperative peer relationships.

## Introduction

Novelty evokes exploratory behaviors in animals ranging from invertebrates to mammals [1–3]. During exploration, animals vary their behavior to sample across the dimensions of a stimulus or environment: hands are moved around to rotate and grip new objects, variable eye movements help view a new scenes, and bodies are moved through the landmarks and edges of a new place. As the structure and values of a stimulus space become learned, behavioral variability decreases, movements become more restricted to sample only the relevant features and can, eventually, become stereotyped habits.

Social novelty can also increase investigative behavior, but the effects of novelty on behavioral variability have not been well studied. By analogy with exploration of space or other stimuli, strangers may initially try-out different behaviors, later converging to particular sets and sequences of interactions as animals learn how best to respond to one-another. There is indirect evidence across animal taxa that interaction variability decreases as relationships form. When fruit flies lose a fight with a particular conspecific, for example, they alter the types of fighting movements used with that individual in the future [4]. Similarly, the formation of dominance relationships in male mice can be observed as an increasingly stable asymmetry of aggressive versus subordinate behaviors within the dyad [5]. These particular examples involve competitive relationships, raising the question of whether social familiarity is also associated with reduced variability in cooperative social relationships. Directly comparing competitive and cooperative relationships can be difficult, in that traditional methods used to investigate relationship formation rely on measures specific to the type of relationship and species studied. It is therefore important to construct metrics that generalize across relationship types, allowing for comparisons between them.

The present study was designed to examine the effects of social novelty on interaction variability in degus (*Octodon degus*), female members of which are predisposed to form and maintain new, cooperative peer relationships. Unrelated female degus nest together and nurse one-another’s young [6–10], and reproductive success is associated with larger, more stable social groups [10,11]. Unrelated male degus are, in contrast, less likely to cohabitate, and reproductive success is not linked to group size [11]. In laboratory settings, female degus interact with strangers more than familiar peers [12], even when they have the option to choose between the two [13]. They also quickly become affiliative with stranger peers in ways that have not been observed in other laboratory rodents [12–14]. Degus therefore offer a unique case study to investigate behavioral patterns associated with the formation of social relationships, one that allows a direct contrast between potentially more competitive relationship types in males and more cooperative in females.

The principal hypothesis of the present work is that animals are more variable in their social interactions when exploring novel conspecifics, and that this holds irrespective of social relationship type. We predict that both male and female degus, with potentially more competitive and cooperative relationships respectively, show decreases in variability as long-term relationships are established. To assess this, we expose same-sex strangers to one-another multiple times over 20 to 40 days and apply measures of interaction variability within and between the reunions. For controls, we also examine the interactions of already-familiar cagemate dyads during the same time period, and subsequently “new cagemates” and “new strangers”. Contrary to expectations, we find that relative variability in strangers versus cagemates are in the opposite direction in female compared with male degus, suggesting that behavioral consistency between encounters depend on the nature of an animal’s relationship. The present study focuses on ethogram-based classification of social interactions; however, one goal of this work has been to establish measures that can be flexibly applied across modes of behavioral observation, species, and setting.

## Results

### Degus interact more with strangers

To help establish validity of the dataset and assess general effects of social novelty, we first tested whether degus interact more with strangers [12,13,15] Degus were separated from a cagemate partner for 24 hours and then united with either a cagemate or stranger in a 50 × 50 cm experimental recording chamber for 20 minutes while video was collected (a “reunion” session, **Figure 1A**). The procedure was repeated 5 times each for cagemates and strangers in an interleaved way, counterbalancing the order between dyads (**Figure 1B**). Consistent with previous findings, degus interacted more with strangers than cagemates (3-factor mixed ANOVA across individuals with exposure day and strangers/cagemate as repeated measures, sex as between-subject; effect of stranger/cagemate: F(1,94)=34.18, p=7.2×10^−8^; **Figure 1C**). Interaction levels were also higher in males across both stranger and cagemate groups (effect of sex: F(1,94)=12.47, p=6.4 × 10^−4^). Interactions were subdivided into five types: agonistic, allogrooming, rear-sniffing, face-to-face, and body-sniffing. There was relatively high variability in the levels and types of interactions both within and across animals over the five cagemate and stranger exposures (**Figure 1D**). However, across the five exposures female degus interacting with strangers showed overall higher levels of agonistic interactions, rear-sniffing, and face-to-face compared against cagemates (2-factor mixed ANOVA, repeated measures over days, false discovery rate correction for 5 tests, agonistic: F(1,51)=8.70, p=0.0048, rear-sniffing: F(1,51)=12.37, p=9.3×10^−4^, and face-to-face: F(1,51)=10.32, p=0.0023; **Figure 1E**, left panels). In males, only agonistic behaviors consistently distinguished strangers and cagemates across exposures (F(1,43)=17.44, p=1.4×10^−4^, **Figure 1E**, right panels).

**Figure 1.**
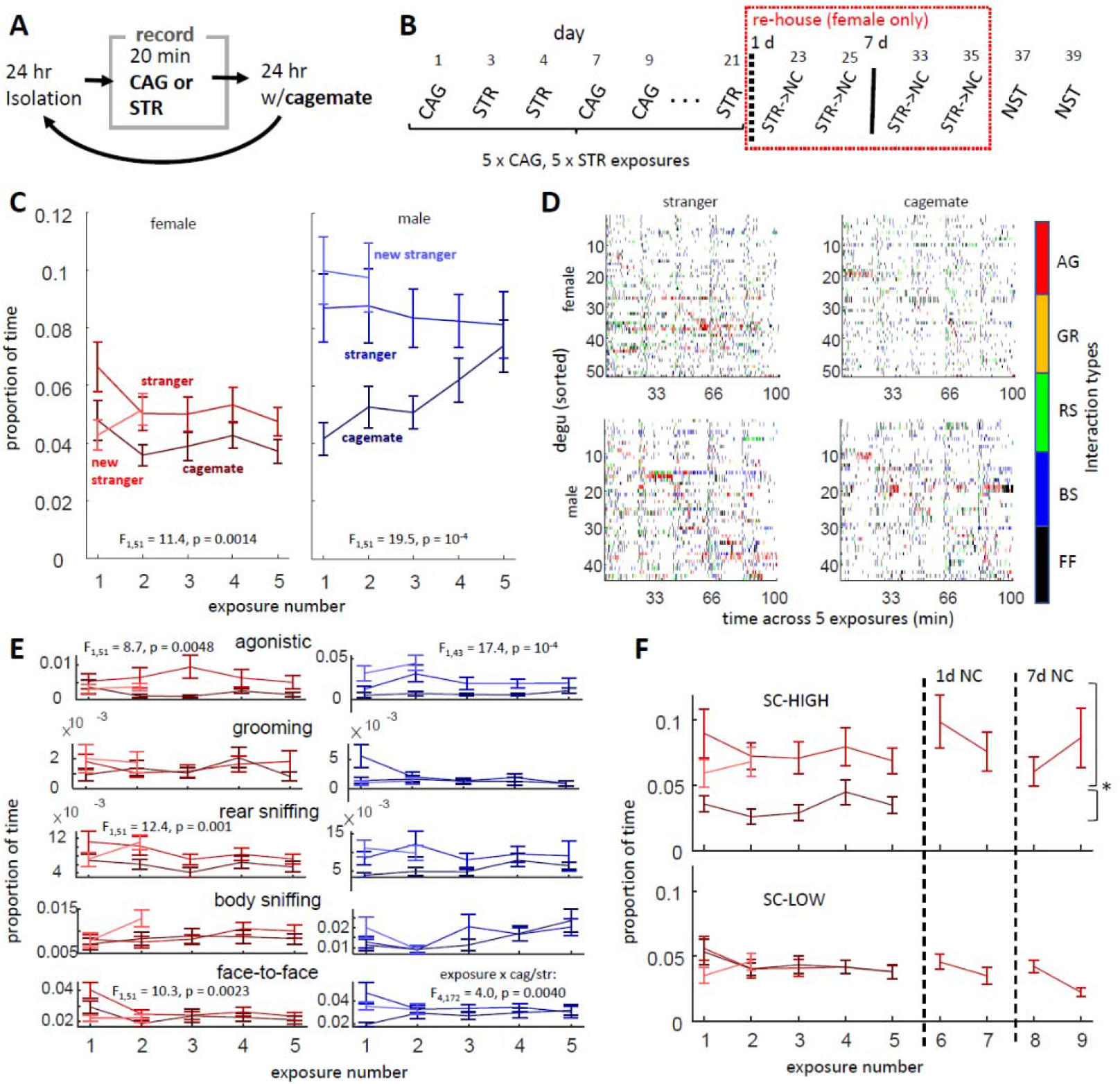
Design and interaction levels. A) Degus received a repeated sequence of 24 hr isolation, 20 minute reunion recording sessions, and 24 hr co-habitation with cagemates. CAG = cagemate reunion, STR = stranger. B) Example sequence of recording sessions. Following interleaved cagemate and stranger reunions, female strangers were co-housed creating “new cagemates” (STR->NC). After all sessions degus were tested with a new stranger (NST). C) Total interaction levels for females (left) and males (right) across the 5 initial exposures (mean ± SEM; statistics are stranger vs. cagemate in 2-way ANOVA). D) Colormaps of interaction types for each degu in 2 s bins across the five exposures. Individual degus (rows) are sorted according to their peak levels of agonistic interactions (AG, red; other abbreviations: GR = allogrooming, RS = rear-sniffing, BS = body sniffing, FF = face-to-face). Higher concentration of face-to-face at session starts helps demarcate the 5 sessions. E) Breakdown of interaction levels across different types of interactions. Higher levels of agonistic, rear-sniffing, and face-to-face were observed in female strangers relative to cagemates (left panels). In males (lower panels), some interaction types decreased and others increased over days. F) Interaction levels over days in female degus that interacted more with strangers (SC-HIGH, top) and those that did not (SC-LOW, bottom). Levels of social interaction between new cagemates (exposures 6 through 9), as well as levels in new strangers (blue), were consistent with those observed in initial strangers on exposures 1 through 5.

We predicted that the difference in interaction levels between strangers and cagemates would diminish as initial strangers became more familiar. This was confirmed by a statistical interaction between stranger/cagemate and exposure day (stranger/cagemate x exposure: F(4,376)=3.16, p=0.014). The pattern was particularly apparent in males, in which face-to-face interactions were more prevalent between strangers during initial exposures (stranger/cagemate x exposure number, false discovery correction for 5 behaviors: F(4,172)=4.00, p=0.0040, **Figure 1E**), with other, non-agonistic behaviors showing similar trends (e.g., rear-sniffing: F(4,172)=2.36, p=0.056). Female degus, in contrast, did not show evidence of diminishing stranger-cagemate differences across days (all interactions: F(4,204)=0.48, p=0.75; individual interaction types: p >>0.05).

To further examine the absence of familiarity effects in females, we subdivided female degus into those that showed significantly higher levels of interaction with strangers (“SC-HIGH”, 16 of 52 degus, determined using a paired t-test on each animal across 5 stranger, 5 cagemate sessions, p < 0.1) and those that did not (“SC-LOW”, 36 of 52 degus). There was no evidence that stranger-cagemate interaction levels changed over days in either the SC-HIGH animal (2-factor repeated-measures ANOVA stranger/cagemate: F(1,15)=83.40, p=1.63×10^−7^, day: F(4,60)=0.89, p=0.47, stranger/cagemate x day: F(4,140)=0.20, p=0.94) or the SC-LOW animals (stranger/cagemate: F(1,35) = 8.0×10^−4^, p=0.98, day: F(4,140)=1.49, p=0.22, stranger/cagemate x day: F(4,140)=0.20, p=0.94; **Figure 1E**). A possible explanation is that the 20 min reunion sessions provided insufficient time for partners to become fully familiarized with one-another. To test this, a subgroup of 40 female degus were co-housed with their stranger partners for 1 day creating new cagemates, then tested in two more reunions, creating “new cagemates”. A smaller subgroup was further co-housed an additional 7 days and tested again twice. Unexpectedly, interactions between SC-HIGH strangers, as new cagemates, remained higher than long-term cagemates following either 1 day (paired t-test, averaging together all cagemate sessions and both new cagemate sessions, t_9_=5.50, p=3.80×10^−4^) or 7 days of co-housing (t_7_=2.98, p=0.021). To investigate whether interaction levels were due to genetic relatedness, we compared sibling versus non-sibling pairs. As only two of the cagemate dyads were non-siblings, this could not be evaluated statistically; however, both members of one, non-sibling cagemate dyad interacted more with strangers (i.e., they were also SC-HIGH animals) while the members of the other dyad did not (were SC-LOW animals). Overall, therefore, female degus exhibited higher interaction levels with strangers than cagemates but there was no evidence these attenuated with familiarity and; consistent with research from other labs [15], this was unlikely to be due exclusively to genetic relatedness.

In males, the differences between interactions with strangers versus cagemates decreased with familiarity, though this pattern could have been influenced by familiarity with the protocol and experimental apparatus. As a final control condition, degus were exposed to a new, unfamiliar conspecific (“new stranger”). In both males and SC-HIGH females, levels of interaction with new strangers were higher than with cagemates (paired t-test between within-animal averages in cagemates and new-strangers, males: t_43_ =5.13, p=6.66×10^−6^, SC-HIGH females: t_15_ =4.17, p= 8.18×10^−4^) and were not lower than interactions with the original strangers (males: t_43_ =1.76, p=0.086 with new stranger > old stranger; SC-HIGH females: t_15_=1.13, p=0.28, **Figure 1F**).

In summary, degus tended to interact more with strangers than with cagemates, though in females this was only the case for one-third of the animals (SC-HIGH group), which continued to interact more even after strangers were co-housed to become new cagemates. Both male degus and SC-HIGH females also interacted more with new strangers than with cagemates.

### In males, strangers have higher session-to-session variability that cagemates; in females, strangers have lower session-to-session variability

The overarching hypothesis of this study is that social interactions are more variable between unfamiliar individuals. Variability can be assessed at multiple timescales, and we began by examining the consistency of a dyad’s interactive behavior between 20 minute reunion sessions. To measure this, we computed interaction vectors: social interaction time budgets across each of 5 interaction types (agonistic, allogrooming, rear-sniffing, body-sniffing, and face-to-face), in each of the two individuals in a dyad. We then computed two, complementary measures: distances (dissimilarity) between interaction vectors, and dyad classification success (the ability of a support vector machine, SVM, to correctly identify which dyad an interaction vector belongs to). While the distance-based method allowed observation of overall variability, the classifier-based method established whether dyads could be distinguished from one-another using combinations of interaction vector elements (**Figures 2A&B**). Notably, the classifier method measures within-dyad variability relative to differences between dyads, which allows specific behaviors to be ignored if inconsistent within a dyad, thereby honing-in on behavior combinations that more selectively define the social relationship.

**Figure 2.**
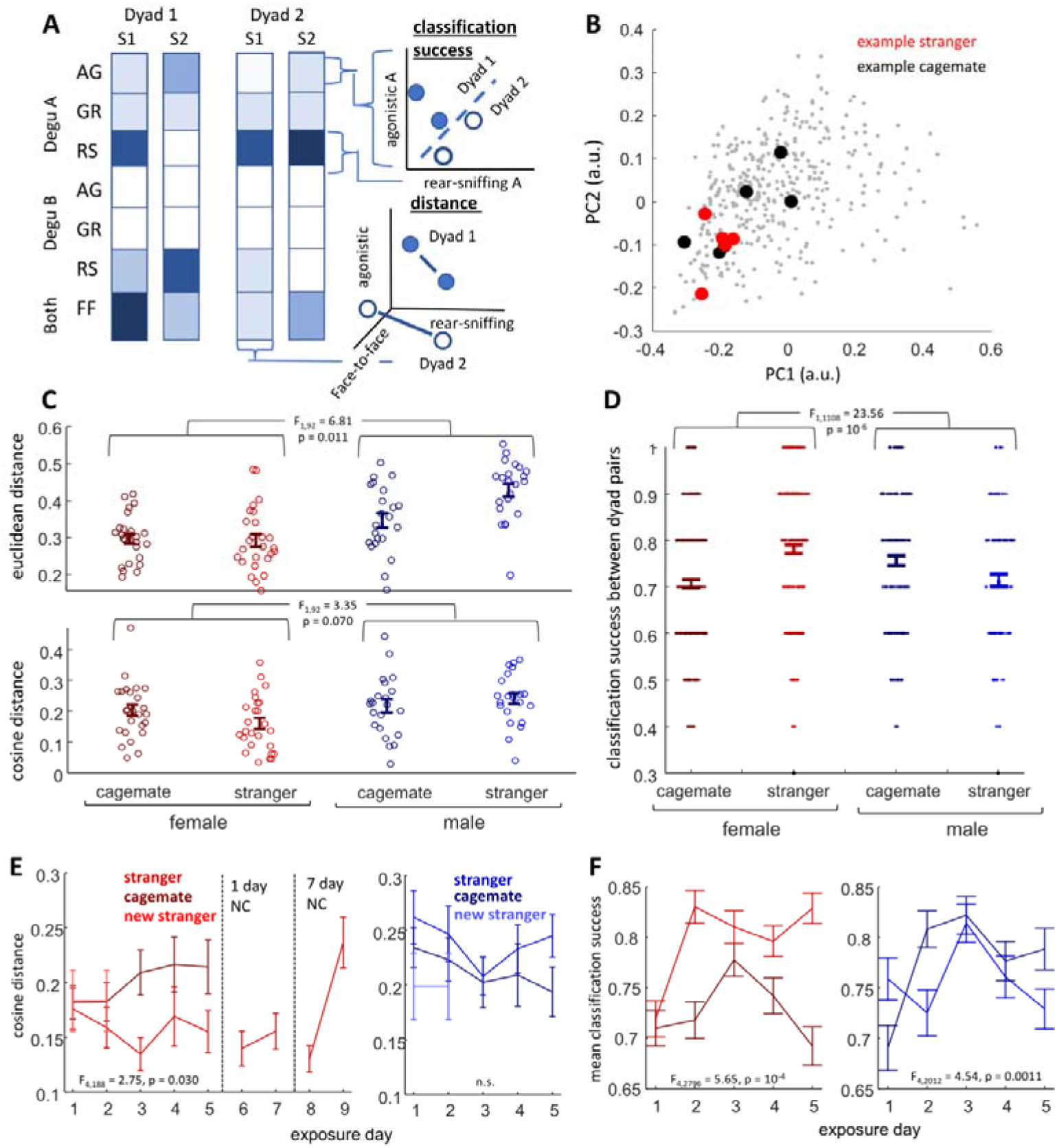
Variability across sessions. A) Diagram illustrating analysis methods. Each column of boxes represents the interaction vector from a single session (abbreviations same as in Figure 1D, body-sniffing omitted for illustration). To the right, vectors are plotted in cartesian coordinates. Top right shows SVM-based classification success, with dashed line representing a hyperplane separating groups in the relevant feature-space. Bottom panel shows interaction vector distances, with points placed in 3 dimensions to emphasize that non-relevant features contribute to the measure. B) Plot of all female dyad interaction vectors using the first two principal components. Dark black dots show all 5 sessions from one example cagemate dyad, red dots from one stranger dyad. In this example, there is higher clustering (less session-to-session variance) in the stranger relative to cagemate. C) Swarm plots showing distribution of within-dyad interaction vector distances (top: Euclidean, bottom: cosine). F- and P-values refer to stranger/cagemate x sex interaction in a 2-way ANOVA. D) Success rate of SVM classifiers discriminating pairs of dyads within each of the four groups (errorbars are mean ± SEM). E) Within-dyad interaction vector distances across exposure numbers (e.g., “exposure 1” only includes distances between exposure 1 interaction vectors and other days). Values are across-dyad averages of within-dyad medians, ±SEM, F- and P-values refer to stranger/cagemate x exposure number interaction using 2-way mixed ANOVA. F) Relative success rates for different days (e.g., “exposure 1” only includes models trained on exposures 2 through 5, then accuracy tested on exposure 1).

Across-session variability was found to differ between cagemates and strangers, but the direction of this difference depended on sex. This could be observed using both interaction vector distances and classifier success rates. Within-dyad distances tended to be lower than between-dyad distances (within-versus between-dyad ratio of 0.73 and 0.69 for females and males respectively), suggesting that individual dyads exhibited unique profiles. Within-dyad, Euclidean distances between interaction vectors were found to be significantly higher in male relative to female degus (2-way ANOVA, effect of sex: F(1,92)=5.5, p=0.021) significantly higher in strangers compared with cagemates (effect of stranger/cagemate: F(1,92)=32.1, p=1.7 × 10^−7^) but relatively higher in male strangers compared with female stranger (stranger/cagemate x sex interaction: F(1,92)=6.8, p=0.011; **Figure 2C, top**). A concern with using Euclidean distance is its sensitivity to total interaction levels; therefore, cosine distance was also used. Cosine interaction vector distances showed differences between males and females (2-way ANOVA, effect of sex: F(1,92)=6.6, p=0.012) and a statistical trend for stranger-cagemate differences to differ between males and females (stranger/cagemate x sex interaction: F(1,92)=3.4, p=0.070; **Figure 2C, bottom**). As described below, trend met criteria for statistical significance when considering exposure number (**Figure 2E**). Contrasting with hypotheses, female strangers appeared to show less variability (more consistency) across sessions relative to female cagemates, male cagemates, or male strangers. This observation females did not appear to be restricted to only dyads with SC-HIGH or only SC-LOW degus (**Supplementary Figure 1A&B**). The interaction between sex and social context depended most on allogrooming and rear-sniffing, as effects decreased when removing either of these components from the interaction vectors, but increased when other interaction types were removed (2-way ANOVA of cosine distances, stranger/cagemate x sex interaction after removing agonistic: F(1,92)=4.3, p=0.040; allogrooming: F(1,92)=2.1, p=0.15; rear-sniffing: F(1,92)=1.65, p=0.20; face-to-face: F(1,92)=4.52, p=0.036; body-sniffing, F(1,92)=6.2, p=0.015; **Supplementary Figure 2A**). Variability differences between males and females appeared to depend more on face-to-face and body-sniffing behaviors (effect of sex after removing agonistic: F(1,92)=7.2, p=0.0085; allogrooming, F(1,92)=8.3, p=0.0050; rear-sniffing, F(1,92)=7.3, p=0.0083; face-to-face: F(1,92)=3.0, p=0.086; body-sniffing: F(1,92)=2.7, p=0.10).

Dyad classification success was evaluated using SVM models trained to discriminate the five, full-session interaction vectors of one dyad from the five interaction vectors of another dyad within the same group (for example, one SVM model might be trained to distinguish one cagemate dyad from another cagemate dyad). Each model was repeatedly trained and tested using a leave-one-out cross validation method. The average success rate was higher than chance, confirming that individual dyads expressed behavioral patterns that were distinguishable from other dyads (binomial probability distribution predicts ∼5% probability of success classifying at least 7 of 10 sessions, mean rates for female cagemates = 7.1, female strangers = 7.8, male cagemates = 7.6, male stranger = 7.3). Rates significantly differed between groups, with classification success higher in female strangers than cagemates, but lower in male strangers than male cagemates (2-way ANOVA effect of stranger-cagemate: F(1,1108)=7.1, p=0.0078, effect of sex: F(1,1108)=0.03, p=0.86, stranger-cagemate x sex interaction: F(1,1108)=23.6, p=1.4×10^−6^; **Figure 2D**). The effects were not due to any one interaction type, as the sex difference remained even after removal of any specific type from the interaction vectors (stranger-cagemate x sex interaction for each: F >20, p <<0.01, **Supplementary Figure 2B**). Higher classification success in strangers was found when considering only female dyads with SC-HIGH degus, and when classifying between dyads with only SC-LOW degus (stranger/cagemate: F(1,313)=6.2, p=0.012, SC-HIGH vs. SC-LOW: F(1,313)=5.4, p=0.020, stranger/cagemate x SC-HIGH/LOW interaction: F(1,313)=0.11, p=0.74; **Supplementary Figure 1C&D**).

We next asked how variability changed over exposures. In females, there was a significant increase in stranger-cagemate differences over days, with variability in stranger dyads decreasing after the first exposure (2-way mixed ANOVA, repeated measures over days, cosine distance: cagemate-stranger: F(1,47)=2.19, p=0.14, exposure day F(4,188)=1.9, p=0.10, cagemate-stranger x exposure day: F(4,188)=2.75, p=0.030); classification success: cagemate-stranger: F(1,699)=29.6, p=7.4×10^−8^, exposure day: F(4,2796)=6.6, p=2.53 × 10^−5^, cagemate-stranger x exposure day: F(4,2796)=5.65, p=1.6×10^−4^ ; **Figures 2E&F, left panels)**. In males, classification success rates appeared to increase in cagemates after the first exposure, while strangers showed less evidence of change (2-way mixed ANOVA, stranger-cagemate: F(1,503)=1.5, p=0.22, exposure day: F(4,2012)=6.1, p=7.74×10^−5^, stranger-cagemate x exposure day: F(4,2012)=4.54, p=0.0011; **Figures 2E&F, right panels**). These patterns were counterintuitive, in that strangers did not converge to become more like cagemates with familiarity; instead, after an initial social exposure in the experimental chamber both female strangers and male cagemates began interacting similarly across days, while female cagemates and male strangers either remained or became less consistent.

### Within-sessions, interaction patterns were not strongly affected by social context

There are no standard measures for variability of social interaction. We selected four measures that provide estimates of variation in the sequencing of actions, the timing between actions, and in blocks of interactions within sessions.

To assess sequencing between actions, we computed transition matrices describing the probability that a “reference” interaction type will be followed by another interaction type. The majority of interactions showed statistically significant relationships (t-test across sessions of probabilities minus chance, baseline probabilities, Benjamini & Hochberg false-discovery correction across 20 comparisons for same-degu interaction pairings and 25 between-degu interaction pairings; **Figure 3A**). The most notable exception was an absence of association between agonistic in one individual with allogrooming in the other. In males, there were also no observable pairings between rear-sniffing and agonistic behaviors across two members of a dyad.

**Figure 3.**
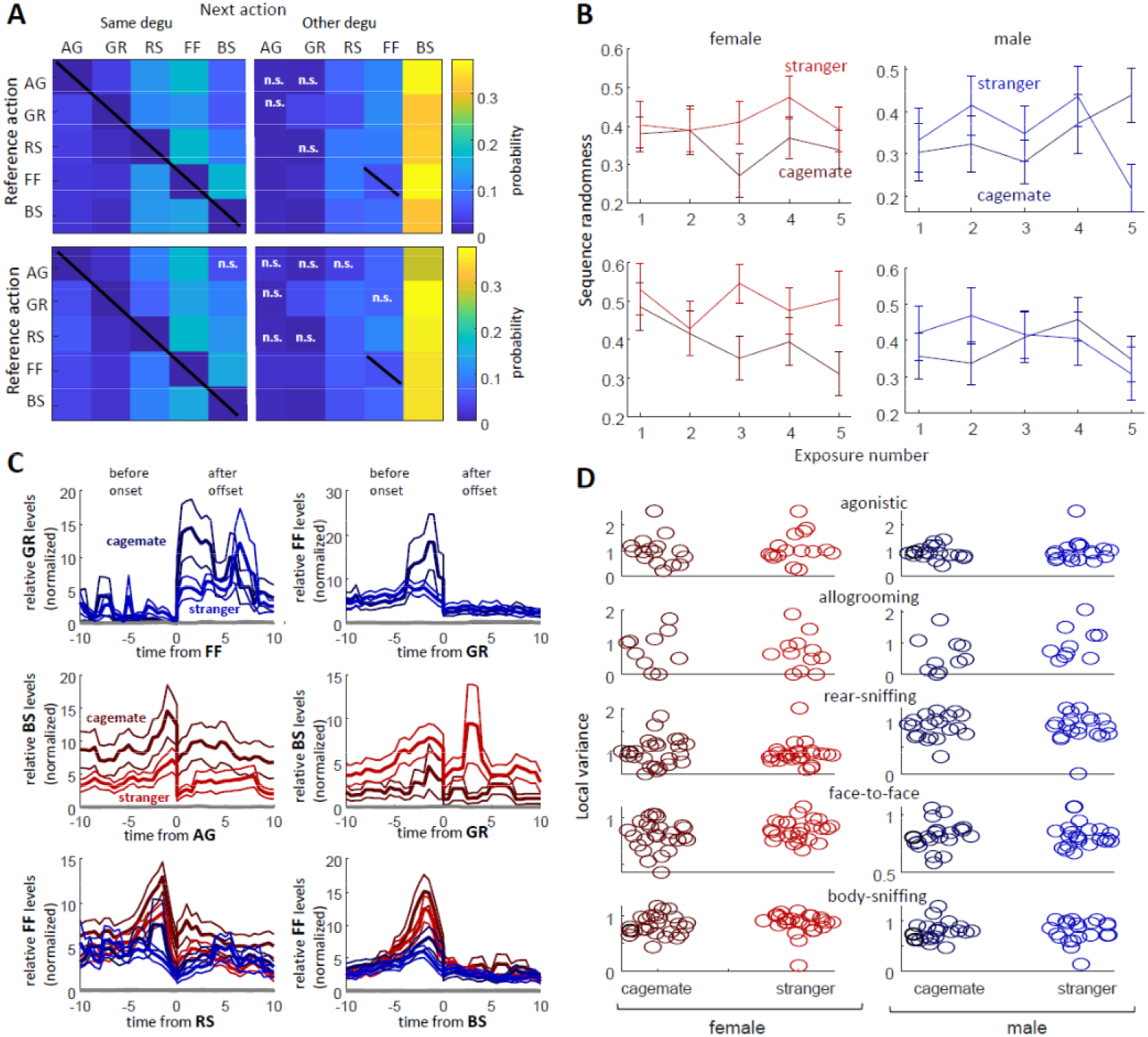
A) Transition matrices between interaction types, averaged across all degus (top: female, bottom: male). Left matrices show probability of the next action (columns) given a reference action (rows) performed by the same degu, right matrices shows probability that the other degu will act, given a reference action in the first degu (matrices computed independently). All cells showed a significant, non-zero probability aside from those labeled “n.s.” Diagonal black lines indicate same-action transitions not included in analysis. Acronyms are the same as in Figure 1D. B) Sequence randomness of interactions across exposures. No significant changes were observed over exposures, nor differences observed between dyad groups in either female (left) or male (right) dyads. C) Example interaction cross-correlograms. The top four emphasize cases of potential difference between cagemates and strangers (thick lines are averages across dyads, thin lines SEM). Top two panels show male levels of allogrooming before and after face-to-face (left) and levels of face-to-face before and after allogrooming (right). Middle panels show female levels of body-sniffing, with cagemates potentially more likely to body-sniff before and after agonistic encounters (left) and less likely than strangers before and after allogrooming (right). Bottom panels show levels of face-to-face interactions around rear-sniffing (left) and body-sniffing (right) in both male (blue) and female (red) dyads. D) Local variance measures examining the consistency of temporal intervals between specific interaction types. No differences between strangers and cagemates were detected.

Although the transition matrices revealed structure in the sequencing of interactions, there was no compelling evidence for overall variability differences between cagemate and stranger groups. This was measured using a sequence randomness score, an index into the overall degree to which behaviors can be predicted given a current behavioral state. Sequence randomness was evaluated across exposures for same-degu and between-degu transition matrices of both sexes. No apparent effects of stranger/cagemate or exposure were observed (mixed 2-way ANOVA, between-subject variability of stranger/cagemate dyad, repeated measures over exposures; female same-degu stranger/cagemate: F(1,4)=0.32, p=0.57, stranger/cagemate x exposure: F(4,128)=0.79, p=0.53; female between-degu, stranger/cagemate: F(1,42)=1.74, p=0.19, stranger/cagemate x exposure: F(4,168)=1.36, p=0.25; **Figure 3B**). When female degus were split into pairs containing SC-HIGH animals and those with only SC-LOW animals, a trend was observed in SC-HIGH for increased, between-degu sequence randomness among strangers (F(1,84)=3.40, p=0.080; **Supplementary Figure 3**). The low effect size and lack of generalizability across sexes, or across all females, emphasized that any differences were very subtle.

Putting aside variability measures, there was also little evidence that stranger and cagemate groups systematically differed in specific interaction transitions (two-sample t-tests, false discovery corrections as above, corrected p-values >0.05). These results did not change when only the first exposure day was considered, nor were differences found in the second-order transition matrix (i.e., the probabilities that one interaction type leads to another, subsequent to the next). Cross-correlations were used to more precisely evaluate the interaction pairings. As with the transition matrices, and as previously observed [12] most pairs of actions showed temporal relationships; however, differences between strangers and cagemates did not exceed threshold for statistical significance (**Supplementary Figure 4**). It is possible that the correction for multiple comparisons occluded subtle differences between cagemate and stranger interaction sequencing. Taking a data exploration (hypothesis-generation) angle, a few specific interaction pairings may have been less likely in strangers: 1) Male strangers may have been less likely to follow face-to-face encounters with allogrooming (two-sample t-test, p=0.020; **Figure 3C**, top-panels), consistent with observations made in females in a prior study (“nuzzling after grooming”, [12]. 2) Male strangers also may have been less likely to precede agonistic behaviors with body-sniffing (p=0.008). 3) female strangers may have been less likely to body-sniff before and after agonistic encounters (p=0.038 and p=0.030), but 3) more likely than cagemates to body-sniff before and after allogrooming (p=0.021 and p=0.025).

Variability of the intervals between interactions of the same type were also analyzed. For example: if face-to-face interactions are a way for two individuals to “check-in” after a period of time had elapsed, these interactions may be more consistently paced in animals who know one-another compared with those who do not. This was measured as the Local Variance of inter-behavior-intervals (IBI’s) for each of the 5 main interaction types There was no evidence for differences in the Local Variance for any type of interaction (two-sample t-tests, false-discovery rate correction, p >> 0.05; **Figure 3D**). The same was true when only the first day of interaction was considered.

A set of final tests examined within-session variability in sets of interactions, by measuring interaction vector distances across blocks of “n” interactions (where n was varied from 1 to 30). As illustrated by **Supplementary Figure 5**, the analysis had fundamental limitations such that distance scores appear higher for sessions with more interactions. When this was controlled for, the data offered no evidence for higher levels of variability in strangers compared with cagemates for either sex.

## Discussion

The present studies were designed to test whether relationship formation involves an initial period of higher variability in the interactions between two individuals. Previous evidence suggests that male and female degus are predisposed to different types of peer relationships, offering an opportunity to compare variability patterns associated with the formation of different types of relationships. We found that interaction variability differed between new and established relationships, but the differences were not consistent between male and female groups. Male stranger dyads showed evidence for higher variability across reunions compared with cagemates—particularly when estimated using dyad classification success. Female stranger dyads, however, were less variable than cagemates across reunions, a reduction that became apparent after either the first or second exposure session when measured using dyad classification success and interaction vector distance respectively. Analyses examining behavior at shorter timescales, within sessions, revealed a structure that is consistent with previous observations [12]; however, only subtle differences were observed between cagemate and stranger groups, with no compelling evidence for higher variability in the latter. As discussed below, degus did appear to interact more with strangers than with cagemates. Together, the results suggest that social exploration may not be expressed as much by behavioral variability, but by higher levels of engagement, in ways that conform to a familiar behavioral structure.

A primary question raised by the results of this study is why, after an initial social exposure, female cagemate dyads exhibited higher levels of session-to-session variability than female strangers. It is possible that the tendency to maintain consistent behavioral patterns helped establish roles in the new relationships. Conversely, it may be that when female degus were placed with a cagemate, they felt less constrained or attentive to the social situation, resulting in behavior that depended more on day-to-day circumstantial factors (e.g., hormonal fluctuations, ambient odors, room temperature, etc.). These explanations are consistent with the notion that being unpredictable is less conducive to forming cooperative relationships. Many primates use specific sets of vocal and physical interactions when encountering strangers, often in stereotyped and ritual sequences. The circumstances in which these behaviors take place—e.g., between similar-ranking males, younger individuals greeting older, and when approaching pregnant females—suggest that the behaviors signal predictability, and function to reduce uncertainty and tension [16–20]. To put another way, reduced variability may help avert high-cost aggressive or competitive interactions. Male degus, unlike females, are not believed to benefit from larger or more stable social groups [11]. Based on the present findings, we hypothesize that a predisposition for more consistent interactions with strangers goes hand-in-hand with a predisposition to form cooperative relationships with new individuals.

One way in which female and male degus were similar is that both interacted more with strangers than with cagemates; however, this increase took-on different patterns. Male strangers showed more greeting (face-to-face) interactions, and a trend for increased investigation (rear-sniffing) during the first few exposures that subsequently attenuated (agonistic interactions with strangers notably became more common after the first social exposure). In female strangers, higher face-to-face, rear-sniffing, and agonistic encounters persisted across reunions (agonistic again being more evident after the first day). Higher levels of interaction with strangers is not surprising, and has been previously observed in our lab and others [12,13,15]. It is also unsurprising given the species’ gregarious behavior in natural habitats: degu burrows are found near one-another, and members of different burrows greet one-another frequently [9,21]. But if female degus form stable, cooperative peer relationships with unrelated individuals, as ecological data suggest [6–10], it is difficult to understand why, in the present study, new cagemates differed from old cagemates. We cannot rule-out the possibility that the structure of the protocol played a role; for example, extended time away from the original cagemates may have sensitized effects of the 24 hour isolation period. It is also possible that degus form fundamentally different bonds with non-siblings encountered in adulthood compared with conspecifics reared together. There are some hints of this from other studies. Females in the wild have been reported to show preferences for members of their own burrow groups during daytime foraging and social greetings [9]. In the lab, genetic kinship can influence social behavior—just not as strongly as familiarity [15]. Future investigation using similar protocols in young animals may help establish whether there are critical periods of development in which certain types of peer bonds take place.

Another notable feature of the present data is that in both males and females, the structure of social behavior was largely preserved between cagemates and strangers. There was no strong evidence for differences in variability in sequencing or temporal relationships of interactions within sessions. Some of the common structure of behavior between groups may be due to the physics of interaction—e.g., two socially attentive degus are more likely to encounter one-another’s faces before body or tail. Or it may be due to the logic of investigative order: the mouth may offer the most important information about another individual, the anogenital region the next most, etc. Additionally, the sequencing and timing of behaviors might be supported by genetic and early-life learning to ensure coordinated activity between individuals of a species. The effects of juvenile social isolation on adult social behavior has been looked at in detail in other species (e.g., chimpanzees: [22,23], rats: [24,25]). While early social deprivation has been shown to have effects on social motivation, the types of interactions used, and an animal’s tendency toward aggression, the sequencing between interactions appears to be relatively resilient to early-life social isolation [25]. Early-life social deprivation may still have subtle effects on the timing of interactions, as well as how vocalizations are used [26], which may make an animal less predictable and thereby destabilizing relationships to greater escalation of aggressive/agonistic encounters [24]. Further work examining interaction sequencing and the effect of early-life isolation in female degus will be valuable for understanding the animals’ abilities to engage constructively with strangers.

Several methodological factors may be critical to interpreting the present results. One of these is the relatively unfamiliar setting of the reunion chamber. Although degus were individually habituated to the chamber over five days prior to the start of the experiment, the room and chamber were still relatively novel, and this may have impacted social behavior between both cagemates and strangers, particularly on the first exposure day. Reactivity of social behavior to environmental novelty has been previously reported in rats [27]. A second, potentially important factor for understanding the results is the relatively few ethogram categories used to track degu interactions. Face-to-face, for example, comprises both nose-to-nose and nose-to-mouth contact, while rear-sniffing includes both anal and genital sniffing. Social interactions can include a high dimensionality of subtle but relevant movements, such as flattening or raising of ears, immobility versus rapid escape, and hesitant versus unrestrained approach. Measuring these additional variables could provide a richer understanding of social dynamics and moment-to-moment metrics of variability. On the other hand, the behaviors that are measured—agonistic (primarily mounting), allogrooming, rear-sniffing, face-to-face, and body-sniffing—are categories that likely have distinct social/ethological relevance for the animals. The low numbers of interactions also simplified the process of identifying which were most responsible for the general patterns. Thus, while the ethogram measures fail to capture the depth of social interaction patterns, they may have helped streamline identification of the similarities and differences between dyads.

While the present project focused its examination on the development of social relationships in degus, an underlying goal was to advance methods that could generalize across species and behavioral measurement tools. In some cases, it may only be possible to interpret an animal’s behavior through its specific ethogram. But while degus are unique in their sociality and peer structures, these same methods could be applied to better understand the formation of cooperative social relationships across species, sexes, and developmental stages. In this way, the study of interaction variability joins a wider movement to apply standardized metrics toward analysis of animal social behavior (e.g., see [28]). As the results demonstrate, however, while distance metrics and classification success can be useful, they cannot be taken in isolation.

## Materials and Methods

### Subjects

Fifty-two adult female degus aged seven to twenty-nine months (mean = 11.9 mo, median = 11 mo) and forty-four male degus aged six to twenty months (mean = 10.4, median = 10) were used. A higher number of female degus were used than males due to a late addition of the “new cagemate” groups in females (see Testing). All animals were housed in same-sex pairs 50.8 × 40.6 × 21.6 cm plastic cages in a breeding vivarium at the University of Montana. Degus were fed a 1:1 mixure of chinchilla and guinea pig “Teklad” feeds (Envigo; Indianapolis, IN). Animals were housed on a 12:12 h light/dark cycle, with all tests occurring during the light (active) cycle. Each cage was also equipped with additional enrichment (e.g., hay, cardboard enclosures, plastic “bones”, and wooden blocks) and, prior to the start of the study, provided with occasional dust baths and handling sessions. Cagemate dyads were created by pairing two same-sex individuals together at the time of weaning. Animals used for stranger dyads were the same as those for cagemates. Pairings were selected with an attempt to keep animals matched for age. Strangers inhabited the same vivarium but in no cases had they been in physical contact.

### Apparatus

All sessions recorded video and audio using a Logitech HD Pro webcam C920 USB2 camera recorded at 30 frames/s. Recording chamber was 50 × 50 × 50 cm painted and sealed wood. Following each social exposure the chamber was cleaned using 70% ethanol then dried.

### Testing

Prior to the reunion procedure, each animal was exposed to the testing chamber without other animals present for at least 5 minutes each day over five days. Twenty-four hours before the first testing session, cagemates were separated into individual cages in the vivarium. Testing began after degus were transported to the testing room and the backside of one degu in each pair was marked for identification (Pet Paint; Camarillo, CA). A pair of either cagemates or strangers was then placed in the chamber and allowed to interact freely for 20 minutes, after which the session ended and animals were returned to home cages in the vivarium with their previous cagemate for 24 hours (**Figure 1A**). By placing cagemates together following each reunion session, animals had the opportunity to “recalibrate” their relationships, preserving it over the weeks of experiment. This cycle of co-housing, isolation, and testing was repeated, with exposures to strangers and cagemates interleaved in a pseudo-randomized order (alternating partners after either one or two reunions with the same partner) for a total of 5 cagemate and 5 stranger sessions (counterbalancing stranger-first and cagemate-first across dyads; **Figure 1B**).

Following 10 reunion sessions, female stranger dyads were co-housed to create “new cagemates”. This was not performed with males to avoid aggression-related injury. Of the 26 stranger female dyads, 24 were made new cagemate and tested after 24 hours of co-housing and 24 hours of isolation, and again after a second 48 hour cycle. A subset of 14 dyads was further tested twice more after 7 days of co-housing. To prevent sampling bias, statistical comparisons that included new-cagemates omitted animals that did not receive this control condition.

Following re-housing reunion sessions in females, and the initial 10 reunion sessions in males, degus were subsequently returned to their original cagemates and then, following 24 hours of isolation, tested with a new stranger. The purpose of the “new stranger” reunions was to establish whether changes in stranger behavior across repeated exposures were due to increased social familiarity, rather than acclimation to the recording chamber or testing protocol.

### Behavioral Scoring

Scoring of physical behavior was performed using BORIS (Behavioral Observation Interactive Research Software; [32]), which allows users to log events during video playback. All analyses were custom-written in MATLAB and are freely available on GitHub.

Filenames for video recordings were changed to ensure raters were blinded to conditions. Raters coded the start and end times of each observed behavior, the type of behavior, and the animal that initiated the behavior. Although the ethogram comprised 17 individual behaviors, for simplicity and consistency with our prior work [12], analyses collapsed behavior types into the following categories: agonistic (mounting, biting, wrestling, boxing, marking, rear-push, tail shaking), allogrooming (sniffing of neck or body with small, repetitive movements), rear-sniffing (anogenital sniffing), face-to-face (nose-to-nose and nose-to-mouth sniffing/contact), and body sniffing (sniffing toward neck and body). Approximately 52% of sessions were scored by author AT and the remainder by 7 undergraduate students all directly trained by AT. Inter-rater reliability (IRR) was estimated using sessions scored both by AT and another student, applying the same transformations and measures used for the primary analyses. Total interaction levels could vary up to a maximum of 20% between raters; however, follow-up examination revealed that inter-rate variance was unlikely to contribute to the primary reported results as 1) sessions were distributed randomly across students and 2) the magnitude of inter-rater error scaled with interaction levels, and analysis controls were used to ensure that interaction levels could not account for interaction variability patterns.

### Measures

All measures were computed from ethogram-based scoring of behavior, performed manually in BORIS. Examination of total interaction time across sessions revealed a tailed distribution that closely resembled a gaussian distribution following a cube root transformation (**Supplementary Figure 7**). To ensure validity of parametric statistical tests and distance metrics, a cube root transform was applied to all “interaction time” measures before performing statistical tests.

#### Interaction time (time budget analysis) allowed comparison with previous datasets and assessment of overall interaction level changes across days

Interaction time was computed as a proportion of session time, and was considered both as a sum total across behavioral types (agonistic, face-to-face, etc.), and separately between behaviors.

#### Between-session variability was estimated using “interaction vector distances” and “dyad classification success”

*Interaction vector distances* were computed by fisrt calculating the proportion of time that each animal spent using a particular type of interactive behavior during a session (an “interaction vector”) and then computing the distance between the vectors associated with different sessions. The types of interactions used for interaction vectors were agonistic, allogrooming, rear-sniffing, body-sniffing, and face-to-face contact. Total vector length was 9 elements, as face-to-face interactions were considered mutual (not assigned to one or the other degu). To compute distances between the interaction vectors, both Euclidean and cosine distance were used. Euclidean distance takes into account total interaction levels, while cosine distance does not. Euclidean and cosine distance are relatively simple, but have advantages over alternatives: e.g., Jensen-Shannon distance (based on Kullback-Leibler divergence) ignores zero elements, which would have misrepresented distances; the Pearson correlation is, like cosine distance, scale invariant, and likely would yield similar values but ignore changes common across all interaction types.

*Dyad classification success* was computed by finding the degree to which a group of interaction vectors could be discriminated from another group. Classification success conflates two variables: within-dyad variability, and between-dyad diversity. The advantage relative to interaction vector distances is that a trained model should ignore types of social interactions that are not useful for discriminating between two dyads, emphasizing those combinations of interactions that are useful. Classification success used a binary support vector machine classifier (SVM, “fitcsvm” in Matlab). SVM works by finding boundaries (i.e., hyperplanes) through n-dimensional space that best differentiate between two sets of data. It’s rate of success in discriminating two sets of data offers a measure of within-versus between-set variability. We used a leave-one-out cross validation method, in which the model is trained on all vectors but one (of 10 interaction vectors, the model is trained on 9), and classification is made on the remaining, repeated for each of the ten interaction vectors across the two dyads. A linear kernel was used in favor of a radial basis function due to ceiling effects in the latter. SVM classifiers were used in favor of neural network classifiers due to the algorithm’s relative tolerance for small training sets.

#### Within-session variability was estimated by examining action-to-action transitions and timing, as well as by comparing interaction vector distances across sets of actions

*Action-to-action transition variability* was computed using “sequence randomness”. Sequence randomness is inversely related to how well an animal’s next action can be predicted, given its previous action. Transition matrices were first computed describing the probability that one type of interaction would lead to one of another type within a 10 s interval (10 s was chosen based on temporal patterns observed in the interaction cross correlograms). Independent transition matrices were computed for interactions performed by the same individual and those performed by the other; for example, if one begins to rear-sniff, the “self” transition matrix estimates the probability that the same animal would subsequently body-sniff, allogroom, etc, while the “other” (between-animal) matrix estimates the probability that the other individual would subsequently perform a body-sniff, allogroom, etc. Rransition matrices did not include probabilities of “no-interaction”. Sequence randomness was the Shannon entropy for each row, averaged across rows:

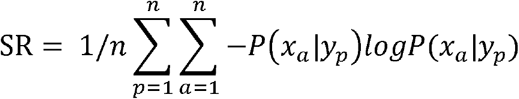

Where x_a_ is interaction type “a”, y_p_ is the immediately preceding interaction type, and “n” is the number of interaction types.

*Variability in action-to-action timing* was estimated using Local Variance and cross correlations. *Local variance* (LV) was used to compute inter-behavior-interval (IBI) variance, a measure commonly used for variance in neuron spike intervals [33]. LV was defined as

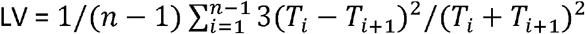

Where T_i_ is the time at which the ith interaction took place and “n” the number of spikes.

Local variance makes no assumptions about the distribution of time intervals between interactions. Empirical evaluation of the IBIs indicate distributions conformed most closely to an exponential (Supplementary Figure 7); i.e., behaviors often come in bursts, similar to what has been described in previous reports [5].

*Action-to-action interactions and timing* were also evaluated using cross correlations (see also [12]). Cross-correlations can be thought of as a histogram where, in this case, times preceding zero (left side of the x-axis) showed the incidence of a target action relative to the initiation time of another, reference action, and times following zero (right side of axis) showed the incidence relative to reference action termination. To make cross-correlations more interpretable, values were normalized by dividing by the proportion of time during the session that a target action took place. Therefore, values of 1 in the cross correlation meant no observable temporal relationship between two interaction types, and y-axis units were normalized rates rather than standard correlation coefficients.

*Within-session, interaction vector distances* were computed using count-based interaction vectors, where each vector element was the sum of interactions of each interaction type over a block of n interactions. If block size n = 10, for example, the vector described how many of 10 consecutive interactions were rear-sniffing events performed by degu A, allogrooming events performed by degu B, face-to-face bouts, etc. This measure helped identify the degree to which dyads changed the set of interactions they were using after a number of behaviors had been performed. Exploratory methods were used to investigate cagemate versus stranger differences across block sizes (1 to 30 interactions per vector), across days. As described in results, many sessions lacked sufficiently high numbers of interactions to yield a single distance value.

### Statistics

Statistical evaluation often involved ANOVA and Student’s t-tests, and only after data were inspected and recognized to approximate gaussian distributions. When assessing interaction levels across exposures, a 2-way repeated measures ANOVA was used (exposure number and cagemate/stranger conditions). When assessing interaction variability changes across exposures, each dyad was considered independent, and mixed models were used (exposure number as a within-dyad repeated measure variable, and cagemate/stranger as a between-subjects variable). In several cases, multiple comparisons had to be taken into account and corrected for. This includes tests that separately examined each interaction type (5 types tested), and when testing significance of temporal relationships between interactions (20 types tested). Significance values are corrected using the Benjamini and Hochberg false discovery rate procedure [36].

### Ethical Note

All protocols were approved by the Institutional Animal Care and Use Committee at University of Montana (protocol number 036-18NIPSYC-061918). Degus were housed with extra space and enrichment and no invasive procedures were performed prior to or during the course of the experiment. Original sample sizes were determined by power analysis performed in G*Power [37] and were based on preliminary effect sizes obtained from within-dyad behavioral patterns across conditions in a prior experiment [12]. Following the experiment, degus were transferred to other experimental protocols; when retired from the laboratory, animals were transferred either to other labs, adopted locally, or humanely euthanized, with post-mortem tissue often used for other research.

## Acknowledgements

Experiment was supported by NIH grant 1R15MH117611-01A1. We would like to thank Stephen Cooke, Kendra Kuehn, and Kendall Butler for help scoring behavior, Kinsey Webb, Janelle Shamp, and Dani Crandell, for help with the degu colony, and Travis Wheeler and Joshua Starmer for initial review of the manuscript and comments on analysis.

## References

1. Berlyne DE. 1966 Curiosity and Exploration. Science 153, 25–33. (doi:10.1126/science.153.3731.25)

2. Hughes RN. 1997 Intrinsic exploration in animals: motives and measurement. Behavioural Processes 41, 213–226. (doi:10.1016/S0376-6357(97)00055-7)

3. Hills TT, Todd PM, Lazer D, Redish AD, Couzin ID. 2015 Exploration versus exploitation in space, mind, and society. Trends in Cognitive Sciences 19, 46–54. (doi:10.1016/j.tics.2014.10.004)

4. Yurkovic A, Wang O, Basu AC, Kravitz EA. 2006 Learning and memory associated with aggression in Drosophila melanogaster. Proceedings of the National Academy of Sciences 103, 17519–17524. (doi:10.1073/pnas.0608211103)

5. Lee W, Fu J, Bouwman N, Farago P, Curley JP. 2019 Temporal microstructure of dyadic social behavior during relationship formation in mice. PLoS ONE 14, e0220596. (doi:10.1371/journal.pone.0220596)

6. Ebensperger LA, Veloso C, Wallem P. 2002 Do female degus communally nest and nurse their pups? Journal of Ethology 20, 143–146. (doi:10.1007/s10164-002-0063-x)

7. Ebensperger LA et al. 2012 Ecological drivers of group living in two populations of the communally rearing rodent, Octodon degus. Behav. Ecol. Sociobiol. (Print) 66, 261–274. (doi:10.1007/s00265-011-1274-3)

8. Ebensperger LA, Villegas Á, Abades S, Hayes LD. 2014 Mean ecological conditions modulate the effects of group living and communal rearing on offspring production and survival. Behavioral Ecology 25, 862–870. (doi:10.1093/beheco/aru061)

9. Ebensperger LA, Hurtado M, Soto-Gamboa M, Lacey EileenA, Chang AnnT. 2004 Communal nesting and kinship in degus (Octodon degus). Naturwissenschaften 91. (doi:10.1007/s00114-004-0545-5)

10. Hayes LD, Chesh AS, Castro RA, Tolhuysen LO, Burger JR, Bhattacharjee J, Ebensperger LA. 2009 Fitness consequences of group living in the degu Octodon degus, a plural breeder rodent with communal care. Animal Behaviour 78, 131–139. (doi:10.1016/j.anbehav.2009.03.022)

11. Ebensperger LA, Correa LA, León C, Ramírez-Estrada J, Abades S, Villegas Á, Hayes LD. 2016 The modulating role of group stability on fitness effects of group size is different in females and males of a communally rearing rodent. J Anim Ecol 85, 1502–1515. (doi:10.1111/1365-2656.12566)

12. Lidhar NK, Thakur A, David A-J, Takehara-Nishiuchi K, Insel N. 2021 Multiple dimensions of social motivation in adult female degus. PLoS ONE 16, e0250219. (doi:10.1371/journal.pone.0250219)

13. Insel N, Shambaugh KL, Beery AK. 2020 Female degus show high sociality but no preference for familiar peers. Behav. Processes 174, 104102. (doi:10.1016/j.beproc.2020.104102)

14. Beery AK, Shambaugh KL. 2021 Comparative Assessment of Familiarity/Novelty Preferences in Rodents. Front. Behav. Neurosci. 15, 648830. (doi:10.3389/fnbeh.2021.648830)

15. Villavicencio CP, Márquez IN, Quispe R, Vásquez RA. 2009 Familiarity and phenotypic similarity influence kin discrimination in the social rodent Octodon degus. Animal Behaviour 78, 377–384. (doi:10.1016/j.anbehav.2009.04.026)

16. Aureli F, Schaffner CM. 2007 Aggression and conflict management at fusion in spider monkeys. Biol. Lett. 3, 147–149. (doi:10.1098/rsbl.2007.0041)

17. Dias PAD, Rodríguez Luna E, Canales Espinosa D. 2008 The functions of the “Greeting Ceremony” among male mantled howlers (Alouatta palliata) on Agaltepec Island, Mexico. American Journal of Primatology 70, 621–628. (doi:10.1002/ajp.20535)

18. Kutsukake N, Suetsugu N, Hasegawa T. 2006 Pattern, Distribution, and Function of Greeting Behavior Among Black-and-White Colobus. Int J Primatol 27, 1271–1291. (doi:10.1007/s10764-006-9072-x)

19. Smuts BB, Watanabe JM. 1990 Social relationships and ritualized greetings in adult male baboons (Papio cynocephalus anubis). International Journal of Primatology 11, 147–172. (doi:10.1007/BF02192786)

20. Whitham JC, Maestripieri D. 2003 Primate Rituals: The Function of Greetings between Male Guinea Baboons. Ethology 109, 847–859. (doi:10.1046/j.0179-1613.2003.00922.x)

21. Fulk GW. 1976 Notes on the Activity, Reproduction, and Social Behavior of Octodon degus. Journal of Mammalogy 57, 495–505. (doi:10.2307/1379298)

22. Kalcher-Sommersguter E, Preuschoft S, Crailsheim K, Franz C. 2011 Social competence of adult chimpanzees (Pan troglodytes) with severe deprivation history: I. An individual approach. Developmental Psychology 47, 77–90. (doi:10.1037/a0020783)

23. Turner CH, Davenport RK, Rogers CM. 1969 The Effect of Early Deprivation on the Social Behavior of Adolescent Chimpanzees. AJP 125, 1531–1536. (doi:10.1176/ajp.125.11.1531)

24. Himmler SM, Himmler BT, Pellis VC, Pellis SM. 2016 Play, variation in play and the development of socially competent rats. Behav 153, 1103–1137. (doi:10.1163/1568539X-00003307)

25. Van Den Berg CL, Van Ree JM, Spruijt BM. 1999 Sequential Analysis of Juvenile Isolation-Induced Decreased Social Behavior in the Adult Rat. Physiology & Behavior 67, 483–488. (doi:10.1016/S0031-9384(99)00062-1)

26. Keesom SM, Finton CJ, Sell GL, Hurley LM. 2017 Early-Life Social Isolation Influences Mouse Ultrasonic Vocalizations during Male-Male Social Encounters. PLoS ONE 12, e0169705. (doi:10.1371/journal.pone.0169705)

27. Varlinskaya E. 2008 Social interactions in adolescent and adult Sprague–Dawley rats: Impact of social deprivation and test context familiarity. Behavioural Brain Research 188, 398–405. (doi:10.1016/j.bbr.2007.11.024)

28. Silk J, Cheney D, Seyfarth R. 2013 A practical guide to the study of social relationships: A Practical Guide to the Study of Social Relationships. Evol. Anthropol. 22, 213–225. (doi:10.1002/evan.21367)

29. Hummer DL, Jechura TJ, Mahoney MM, Lee TM. 2007 Gonadal hormone effects on entrained and free-running circadian activity rhythms in the developing diurnal rodent Octodon degus. Am J Physiol Regul Integr Comp Physiol 292, R586–597. (doi:10.1152/ajpregu.00043.2006)

30. Mahoney MM, Rossi BV, Hagenauer MH, Lee TM. 2011 Characterization of the Estrous Cycle in Octodon degus. Biology of Reproduction 84, 664–671. (doi:10.1095/biolreprod.110.087403)

31. Lee TM. 2004 Octodon degus: a diurnal, social, and long-lived rodent. ILAR J 45, 14–24. (doi:10.1093/ilar.45.1.14)

32. Friard O, Gamba M. 2016 BORISIZ: a free, versatile open-source event-logging software for video/audio coding and live observations. Methods Ecol Evol 7, 1325–1330. (doi:10.1111/2041-210X.12584)

33. Shinomoto S, Miura K, Koyama S. 2005 A measure of local variation of inter-spike intervals. Biosystems 79, 67–72. (doi:10.1016/j.biosystems.2004.09.023)

34. Benjamini Y, Hochberg Y. 1995 Controlling the False Discovery Rate: A Practical and Powerful Approach to Multiple Testing. Journal of the Royal Statistical Society: Series B (Methodological) 57, 289–300. (doi:10.1111/j.2517-6161.1995.tb02031.x)

35. Faul F, Erdfelder E, Lang A-G, Buchner A. 2007 G*Power 3: A flexible statistical power analysis program for the social, behavioral, and biomedical sciences. Behavior Research Methods 39, 175–191. (doi:10.3758/BF03193146)

