## Supplementary Figures for "New relationships are not necessarily more variable: female degus are more consistent with strangers, male degus are less"

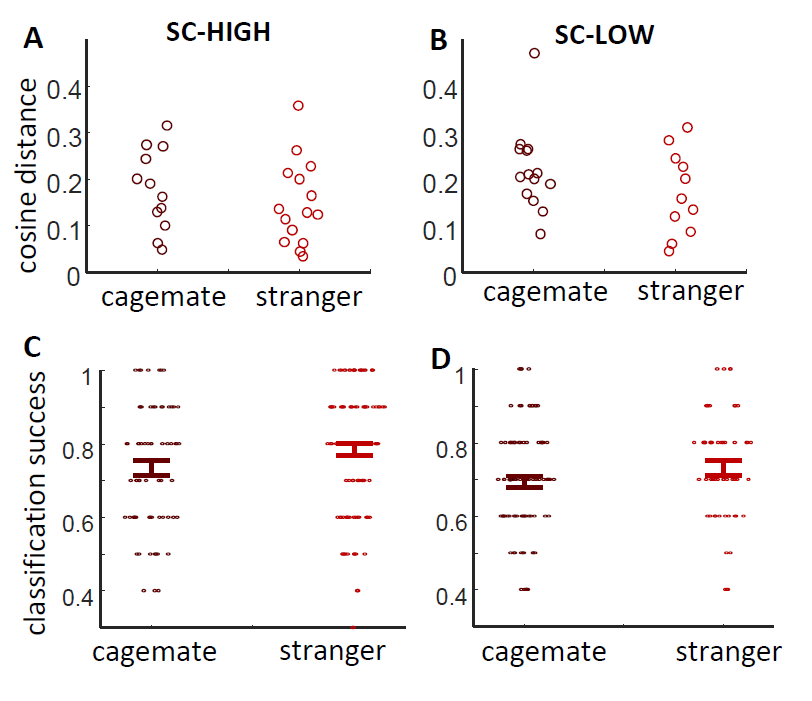

**Supplementary Figure 1:** Variability comparison between cagemate and stranger dyads in SC-HIGH versus SC-LOW females. A & B) Swarm plot showing distribution of cosine distances in cagemate and stranger dyads in SC-HIGH (A) and SC-LOW (B) dyad groups. In this case, all exposure days are combined, limiting detection of cagemate-stranger differences; however, for both SC-HIGH and SC-LOW dyad groups, mean values were lower in stranger (2-way ANOVA effect of stranger/cagemate: F(1,48) = 2.6, p = 0.12; effect of SC-HIGH/LOW: F(1,48) = 1.7, p = 0.20; stranger/cagemate x SC-HIGH/LOW interaction: F(1,48) = 0.17, p = 0.68). C & D) Overlaid errorbar and swarm plots showing relative classification success rats in stranger and cagemate dyads in SC-HIGH (C) and SC-LOW (B) female groups. The classifier showed moderately higher success rates in the SC-HIGH group; however, there was no indication that this impacted the stranger-cagemate differences (2-way ANOVA effect of stranger/cagemate: F(1,313) = 5.43, p = 0.020; effect of SC-HIGH/LOW: F(1,313) = 6.5, p = 0.012, stranger/cagemate x SC-HIGH/LOW interaction: F(1,313) = 0.11, p = 0.74). Together the results suggest that the increased levels of interaction with strangers observed in SC-HIGH group did not impact levels of session-to-session variability.

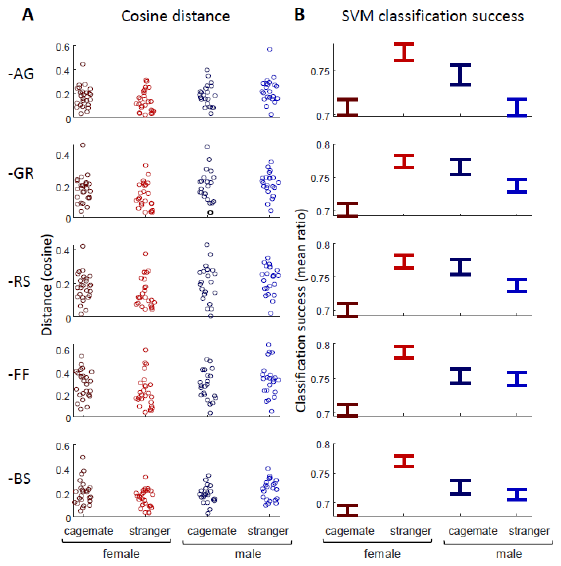

**Supplementary Figure 1: Social familiarity x Sex effects are preserved in reduced interaction vectors.** A) Within-dyad cosine distances across cagemate/stranger and male/female groups after removing individual interaction types from the interaction vector (-AG = agonistic removed, GR = allogrooming, RS = rear-sniffing, FF = face-to-face, BS = body-sniffing). Female stranger dyads consistently showed lower variance than other groups, though the effect of cagemate/stranger x sex interaction became weak when grooming and rear-sniffing were removed. B) Same as part A but for dyad classification success. Female strangers were consistently easier to classify than female cagemates, while classification success in male strangers was typically lower than cagemates.

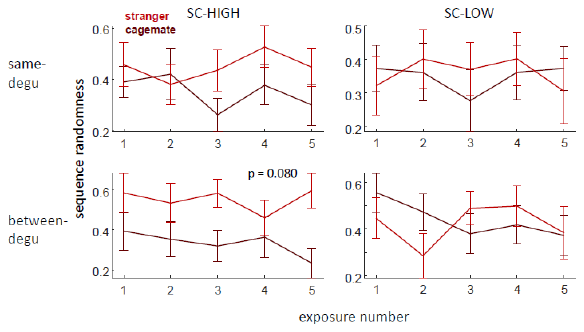

**Supplementary Figure 3: Interaction predictability (sequence randomness) in SC-HIGH and SC-LOW females.** Each plot shows “sequence randomness” scores across exposures in stranger and cagemate females; a measure of the predictability of one social interaction, given an immediately preceding interaction. Top panels show sequence randomness for interactions performed by the same degu (i.e., the same degu performed one interactive behavior after another), bottom panels show scores for interactions from one degu to another (often degus response to the other’s action). A statistical trend suggested higher, between-degu sequence randomness across exposures in the SC-HIGH but not SC-LOW group; however, differences were not statistically significant and subject to multiple comparisons.

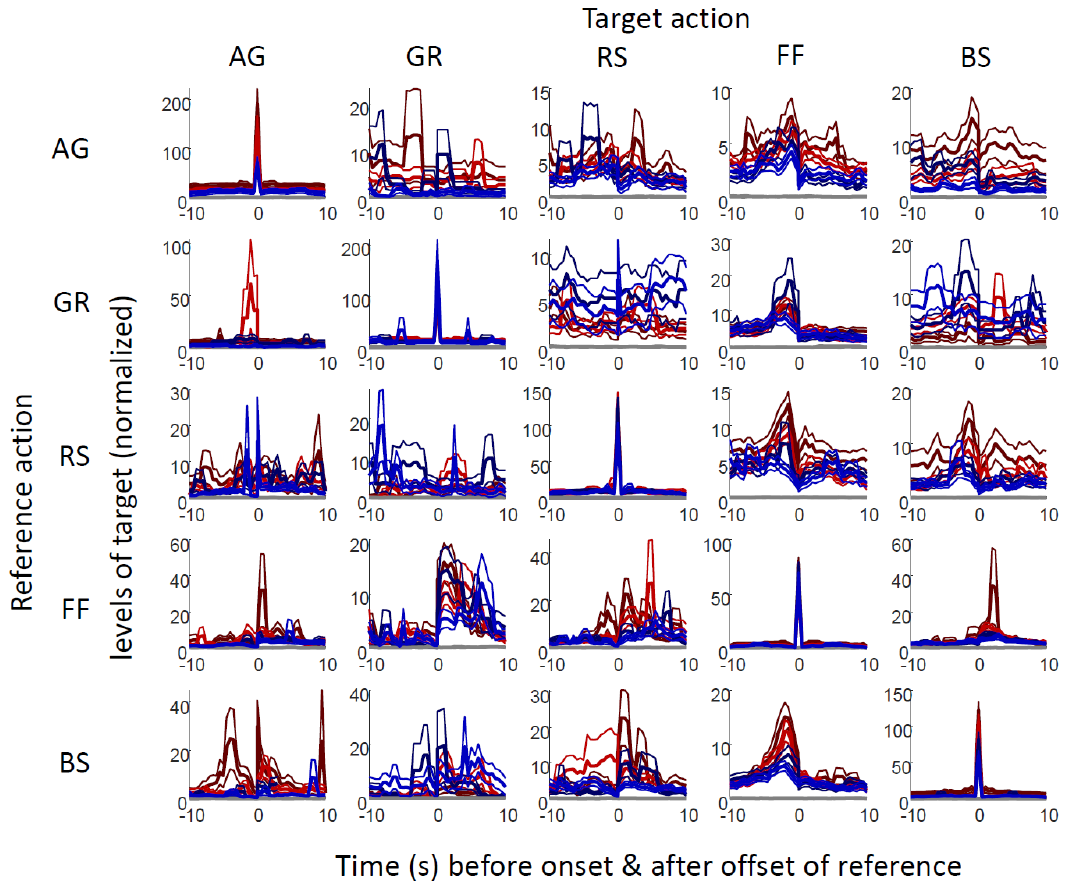

**Supplementary Figure 4: Cross correlograms between all interaction types.** Levels of a “target” interaction preceding onset or following the offset of a “reference” interaction are plotted for each of the 5 examined interaction types (AG = agonistic, GR = allogrooming, RS = rear-sniffing, FF = face-to-face, BS = body-sniffing). Red traces are averages in females (light red = stranger, dark red = cagemate) and blue males (light blue = stranger, dark blue = cagemate). Strong asymmetries were observed similar to those reported in previous work; e.g., face-to-face interactions—often greetings—consistently preceded other types of interactions (illustrated by the left asymmetry in the 4^th^ “FF” column). Due to variance across sessions and between dyads, statistically significant differences between cagemates and strangers were rare, though examples of potential differences are described in the main text.

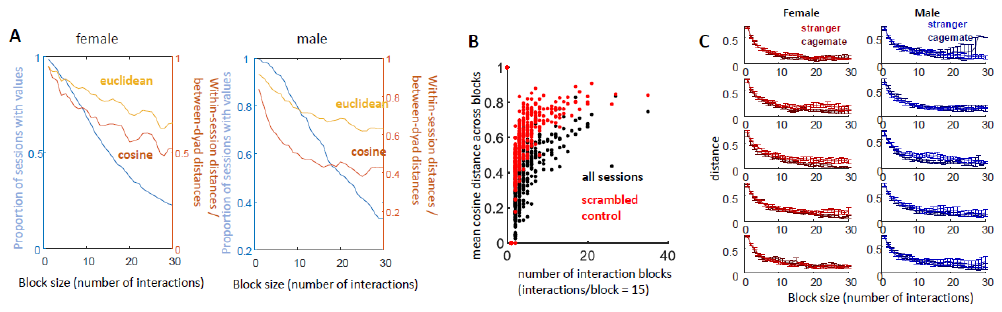

**Supplementary Figure 5: Within-session variability measured by distance between interaction blocks.** As with between-session analyses, variability was measured as interaction vector distance, in this case computed as the number of interactions of each type taking place over a block of “n” interactions, where n was varied from 1 to 30. **A)** An important caveat to the analysis is that is that a reunion session only has a distance value if there are at least two blocks; thus, as the block size (n) is increased, the number of sessions included in the analysis decreases (in the plots, this is shown by the light-blue trace). In female degus (left plot), the number of sessions decreased to below 50% when n = 16 (i.e., fewer than 50% of the sessions included more than 32 interactions); in males (right plot), 50% was reached when n = 22. Using cosine distance (dark orange trace), the ratio of within- versus between-dyad distances reached a minimum of 0.48 in females (0.65 at n = 15) and 0.38 in males (0.45 at n = 21). Euclidean distances were associated with higher ratios of distances within versus between-dyads (light orange traces). **B)** Another caveat with the analysis is that results can depend on how multiple blocks within a session are handled. If averaging across all distances within a session, values become inflated within increasing numbers of blocks in a session. This is illustrated here for block sizes of 15 interactions. Black points are all reunion sessions plotted according to number of 15-interaction blocks (x-axis) and mean cosine distance. This effect is further increased after interaction identity is scrambled (red points) and can be attributed to the accumulation of variance with increasing blocks. **C)** Averaged, within-session distances for each block size (x-axis) across the 5 exposures sessions (top to bottom panels) in females (left panels) and males (right panels). Across multiple block sizes and days strangers showed higher distances (higher variability; estimated using a regression model that included block size, exposure session, and cagemate/stranger category). However, these differences disappeared when using methods that controlled for the scaling of distances with session block number.

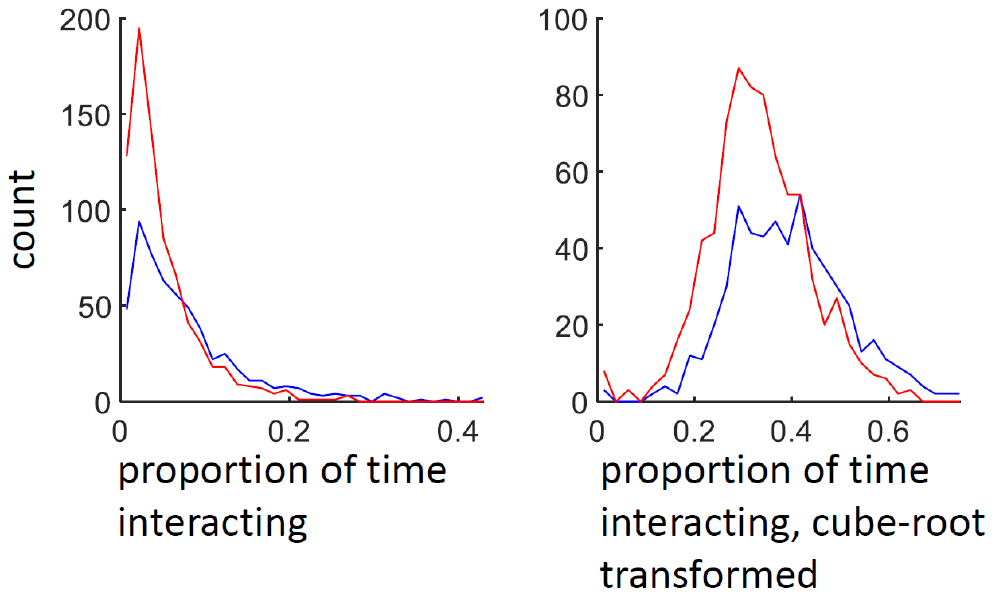

**Supplementary Figure 6: Histograms of interaction time.** Left: Actual time spent interacting. Right: Cube-root transformed interaction time. Interaction time was measured as a proportion of total session time, to account for variations in total time of 20 minutes. The transformation aiding in analysis and interpretation.

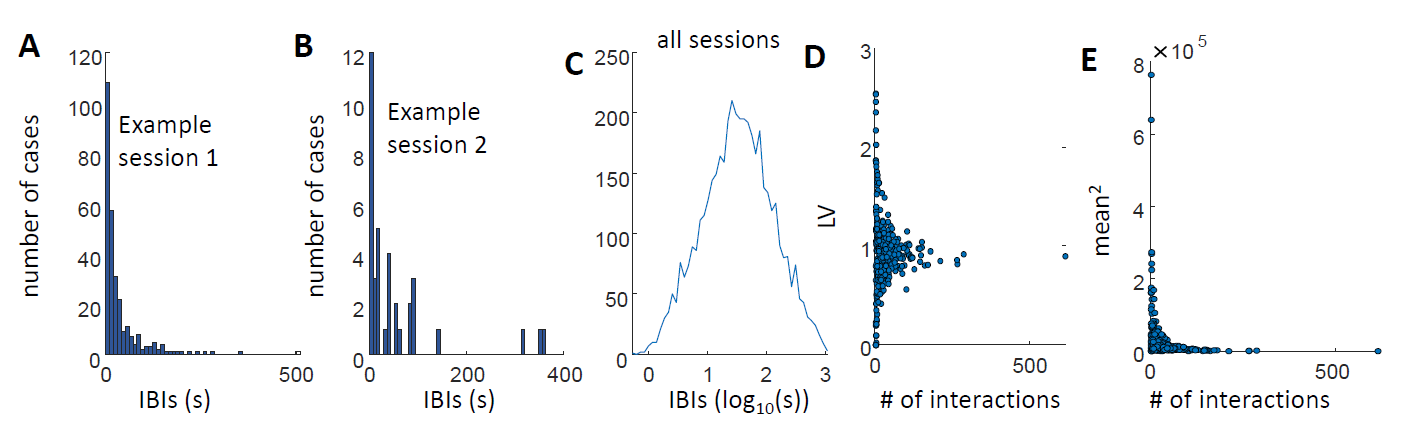

**Supplementary Figure 7: Illustration of inter-behavior-intervals**. A) Histogram of the intervals between face-to-face interactions in one example session. Distribution suggests the interval vary exponentially. B) Histogram of intervals between face-to-face interactions in another example session with far fewer interactions. Decreasing the number of interactions does not strongly impact the exponential fit of the curve. C) All inter- face-to-face intervals plotted on a log scale. Approximate gaussian distribution suggests that a log-normal fit. D) Local variance measure applied to all interaction types, across all sessions reveals that higher numbers of interactions results in smaller deviations from LV = 1. E) When computing the mean-squared of the inter-behavior intervals—the variance of the exponential distribution—a very strong relationship between number of interactions and variance is observed.
